# The sequence and structural integrity of the SARS-CoV-2 Spike protein transmembrane domain is crucial for viral entry

**DOI:** 10.1101/2024.11.19.624333

**Authors:** J Ortiz-Mateu, D Belda, AI Avilés-Alía, J Alonso-Romero, MJ García-Murria, I Mingarro, R Geller, L Martinez-Gil

## Abstract

The Spike (S) protein of SARS-CoV-2 is a type I membrane protein that mediates target cell recognition and membrane fusion. While its transmembrane domain (TMD) is traditionally viewed as a passive anchor to the viral envelope, emerging evidence suggests that TMDs often play active roles in the biogenesis and function of membrane proteins. Here, we investigated the functional role of the SARS-CoV-2 S protein TMD during viral entry. To this end, we introduced a series of amino acid substitutions and insertions within the hydrophobic core of the TMD and assessed their impact on S protein activity. Our findings reveal that the SARS-CoV-2 S protein is susceptible to alterations in its TMD. Functional determinants, including sequence features and structural parameters critical for viral entry, are distributed throughout the TMD, with a more pronounced contribution from its N-terminal region. We also demonstrate that the relative orientation of the regions flanking the TMD influences viral entry. Finally, our data suggest that the TMD mediates homo-oligomerization through a motif enriched in small residues, underscoring its functional importance beyond membrane anchoring.

## Introduction

It is becoming increasingly clear that a deep understanding of the molecular mechanisms occurring during viral infection is required for an effective rational design of antiviral agents and vaccine immunogens. In the case of SARS-CoV-2, special attention has been given to the entry process and the role that the Spike (S) protein plays in it^1^. The SARS-CoV-2 S protein is a trimeric class I fusion glycoprotein that mediates target cell recognition and membrane fusion^2,3,4^. During post-translational maturation, the full-length protein is cleaved into two subunits, S1 and S2. S1 contains the receptor-binding domain (RBD) and facilitates attachment to the angiotensin-converting enzyme 2 (ACE2) on the host cell surface. S2 drives the fusion of the viral and host membranes and encompasses the fusion peptide, two heptad repeat regions, and a C-terminal transmembrane domain (TMD).

Often, the TMD is seen as a passive physical anchor whose sole function is to attach the protein to the viral envelope. However, current knowledge of the structure, dynamics, and biogenesis of integral membrane proteins demonstrates that TMDs, alongside membrane-proximal regions, play key functions beyond membrane anchoring^5–7^. Furthermore, in recent years, significant advances in understanding the role of the TMDs of viral fusion proteins have been made highlighting their critical role in the fusion process, and also demonstrating their involvement in several aspects of the viral lifecycle^8,9^.

Broer et al. demonstrated that substituting the N-terminal membrane proximal region, the TMD, and the endodomain of the SARS-CoV-1 S protein with the corresponding domains of the vesicular stomatitis virus (VSV) Glycoprotein greatly affects the protein’s function^10^. A few years later, a mutagenesis-based study, once again using VSV pseudo-particles, revealed that substituting Gly_1201_, Ala_1204_, Val_1210_, and Leu_1216_ by Lys residues in the core hydrophobic region of the SARS-CoV-1 S protein resulted in reduced infectivity^11^ likely by altering the hydrophobic nature of the TMD.

The SARS-CoV-2 S protein TMD has been linked with trimerization. A chimera including the RBD followed by the TMD was identified as a trimer in which the RBD was properly folded, suggesting that the TMD acts as an oligomerization domain^12^. However, there is some controversy regarding the oligomerization motif within the TMD. Abery et al. identified a GxxxG motif (a sequence known to promote oligomerization of TMDs^13^) within the TMD of the SARS-CoV-1 S protein and demonstrated its importance in the trimerization of the protein^14^. Shortly after, Corver et al.’s results indicated that these residues were not important for the trimerization of the protein or viral entry^15^. Recently, it has been suggested that a new motif based on a hydrophobic zipper mediates the trimerization of SARS-CoV-2 S protein TMD^16^.

Several structures of the S protein have been solved^17–19^. However, only one includes the TMD^20^. In this structure, the S protein was found in a post-fusion state where the fusion peptide (FP) forms a hairpin-like wedge spanning the entire lipid bilayer and the TMD wraps around it. Two contact points between the FP and the TMD were described involving Phe_1220_ and Leu_1234_ on the TMD.

Here, we explore the role of the SARS-CoV-2 S protein TMD during viral entry. To do so we generate a series of variations within the hydrophobic core of the TMD and analyze their impact on the protein function. Furthermore, we investigate the potential of the TMD to induce oligomerization as well as the structural and sequence parameters responsible for it. Our results demonstrate that the SARS-CoV-2 S protein TMD plays a key role in the protein function beyond membrane anchoring. The sequence and structural parameters of the TMD relevant for viral entry are distributed across the entire segment, which might explain its strong sequence conservation and the literature controversy. Additionally, our results suggest that the relative orientation of the regions before and after the TMD is important for particle entry. Finally, we identify that the SARS-CoV-2 S protein TMD is capable of forming homo-oligomers and these occur through a motif in which small residues such as glycine and alanine are relevant, results that were corroborated *in silico*.

## Results

### Pseudotyped-VSV particles for the analysis of viral entry

The SARS-CoV-2 S protein contains a single TMD in the C-terminal region of the protein (Fig. 1A). It spans 23 amino acids from Trp_1214_ to Cys_1236_ with a predicted ΔG_app_ of -3.4 kcal/mol for its insertion into the membrane, according to the Δ*G Prediction Server*^21,22^. The predicted TMD includes an aromatic-rich region (Trp_1214_-Tyr-Ile-Trp_1217_) in its N-terminal end and a Cys pair (Cys_1235_-Cys_1236_) in its C-terminal end. However, the structure of the S trimer in its post-fusion state^20^ shows a helical secondary structure from Leu_1218_ to Leu_1234_. Although the aromatic region and the C-terminal Cys-rich section on the TMD most likely interact with the membrane either as an interfacial membrane proximal region^20,23^ or through palmitoyl post-translational modifications^24–27^, in the present study we focus exclusively on the helical hydrophobic core of the TMD, that is, from Leu_1218_ to Leu_1234_. The sequence of all the mutations included in this paper and their insertion potential can be found in Fig.1B.

**Figure 1.**
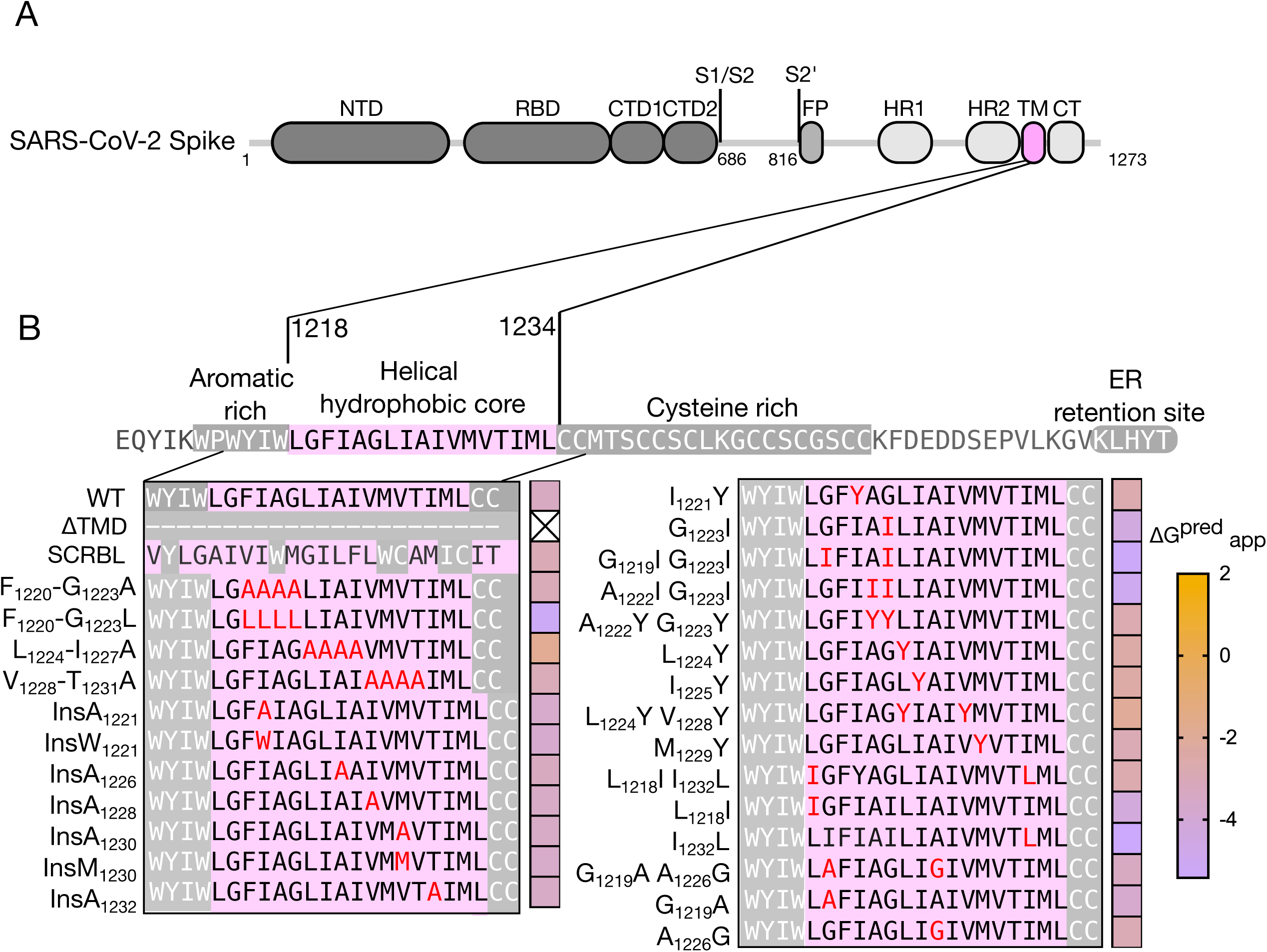
Schematic representation of the SARS-CoV-2 spike protein and its TMD. **A**. Schematic representation of SARS-CoV-2 Spike protein domain architecture. The position of key domains is highlighted: NTD, N-terminal domain; RBD, Receptor Binding Domain; CTD1, C-terminal domain 1; CTD2, C-terminal domain 2; S1/S2, S1/S2 cleavage site; S2′, S2′ cleavage site; FP, fusion peptide; HR1, heptad repeat 1; HR2, heptad repeat 2; TM, transmembrane domain; CT, cytoplasmic tail. **B**. Sequence of the SARS-CoV-2 S protein TMD and the mutations included in this study. The sequence of the SARS-CoV-2 S protein from Glu_1206_ to Thr_1273_ is shown. Membrane-associated regions, including an aromatic-rich section and a cysteine-rich domain, are highlighted with light letters on a gray background. The TMD hydrophobic core is shown with dark letters on a pink background. The TMD, as predicted by the ΔG Prediction Server (WT), and the variations included in this work are highlighted. A heat map (right of the sequence) shows the predicted ΔG_app_ for each TMD mutant.

To study the role of the SARS-CoV-2 S protein TMD during viral entry we utilized pseudotyped vesicular stomatitis virus (VSV) particles coated with different S protein mutants (Fig. 2A)^10,15,28,29^. Briefly, HEK293T cells were transfected with plasmids encoding the different S mutants and subsequently infected with a recombinant VSV in which the glycoprotein gene has been replaced with GFP (VSVΔG-GFP). Next, the infectivity of the resulting particles was assayed on VeroE6 cells expressing the TMPRSS2 cofactor by counting GFP-positive cells. VSVΔG-GFP particles were pseudotyped with the native SARS-CoV-2 Wuhan-Hu-1 S protein as a positive control and reference value (WT). No infection was observed when the TMD of the SARS-CoV-2 S protein was eliminated (ΔTMD) or was substituted by a scrambled version of the TMD (SCRBL) where the composition of the residues included in the TMD was preserved but their position was randomly varied, confirming the importance of the SARS-CoV-2 S protein TMD in the entry process and indicating a role beyond membrane anchoring (Fig 2B). Expression of the chimeras included in this work can be found in Supp Fig 1.

**Figure 2.**
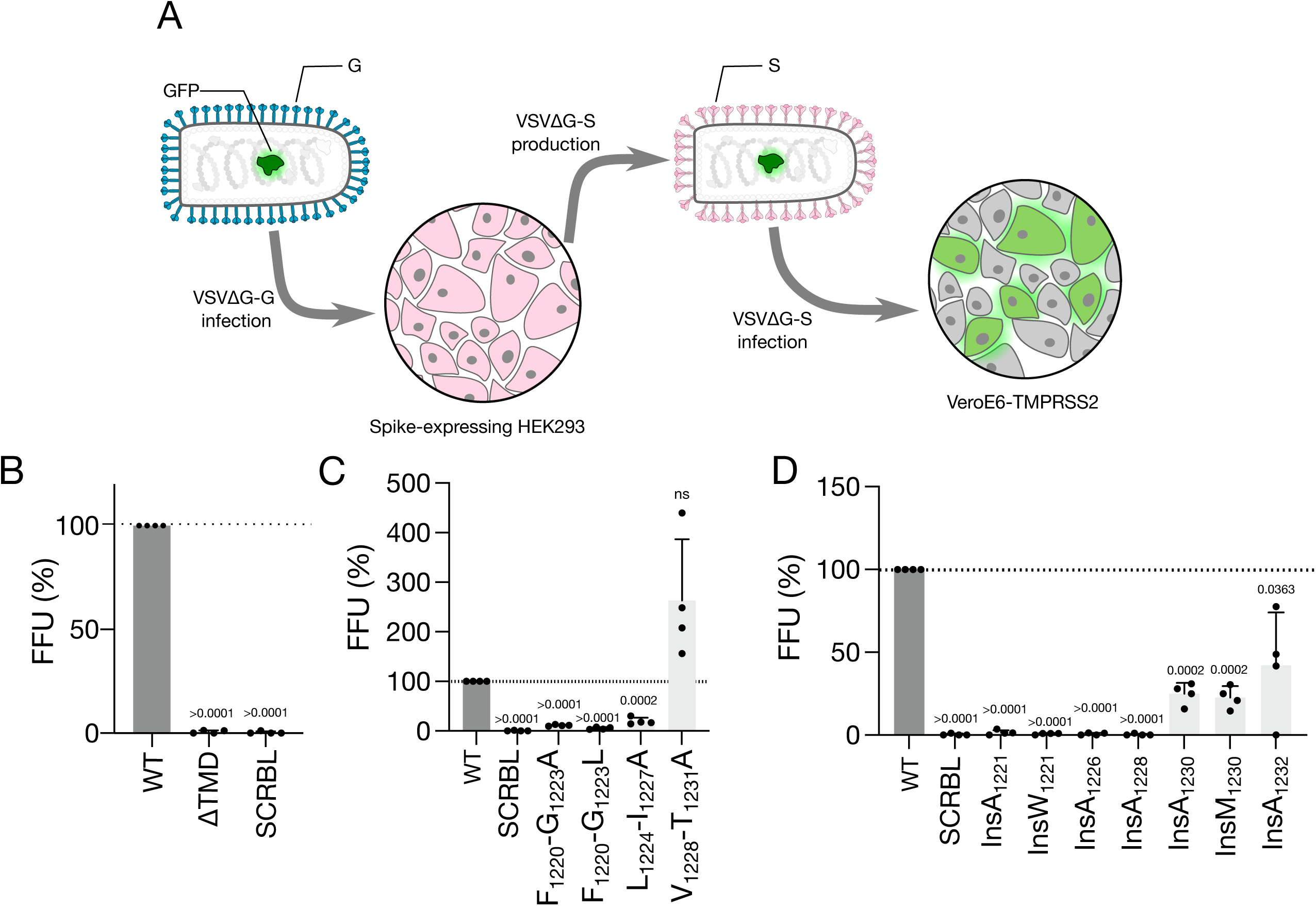
VSVΔG-GFP pseudotyped virus assay. **A**. Generation of VSVΔG-GFP pseudotyped particles. HEK293T cells were transfected with the corresponding SARS-CoV-2 S protein mutant and infected with a VSV lacking the G gene and expressing GFP (VSVΔG-GFP). The resulting virus, incorporating the S mutants on its surface (VSVΔG-GFP -S), was recovered from the media and used to infect Vero-TMPRSS2 cells. Viral infectivity was then analyzed by counting GFP-positive cells. **B-D**. Focus Forming Units (FFU) measured after infection with VSVΔG-GFP pseudotyped with different mutants of the SARS-CoV-2 S protein TMD. All samples were normalized to the wild-type SARS-CoV-2 S protein (WT). The p-values (<0.05) for individual one-sample t-tests (vs WT) are indicated above each bar. **B**. FFU from VSVΔG-GFP particles decorated with the wild-type SARS-CoV-2 S protein (WT), the SARS-CoV-2 S protein where TMD was eliminated (ΔTMD), and a scrambled version of the SARS-CoV-2 S protein TMD (SCRBL). **C**. FFU from pseudotyped VSVΔG-GFP with the SARS-CoV-2 S protein with substitutions of amino acid stretches, Phe_1220_ to Gly_1223_, Leu_1224_ to Ile_1227_, and Val_1228_ to Thr_1231_, with alanine (Phe_1220_-Gly_1223_A, Leu_1224_-Ile_1227_A, and Val_1228_-Thr_1231_A respectively). Substitutions of amino acids Phe_1220_ to Gly_1223_ to leucine (Phe_1220_ to Gly_1223_L) were also included. **D**. FFU from pseudotyped VSVΔG-GFP with the SARS-CoV-2 S protein including alanine insertions at positions 1221, 1226, 1228, 1230, and 1232 (InsA_1221_, InsA_1226_, InsA_1228_, InsA_1230_, and InsA_1232_), tryptophan insertion at position 1221 (InsW_1221_), and methionine insertion at position 1230 (InsM_1230_).

Next, stretches of four amino acids (one helical turn) on the SARS-CoV-2 S protein TMD, starting at Phe_1220_, were substituted with alanine residues (Fig 2C). Substitution of the section covering Phe_1220_-Gly_1223_ and Leu_1224_-Ile_1227_ (F_1220_-G_1223_A and L_1224_-I_1227_A respectively) greatly diminishes the infectivity of the pseudo-typed VSVΔG-GFP particles. Substitution of Val_1228_-Thr_1231_ (V_1228_-T_1231_A) to alanines did not decrease viral entry, on the contrary, a non-significant increase was observed. To ensure that the observed results are not biased by substitutions to Ala we substitute the Phe_1220_-Gly_1223_ with Leu residues (F_1220_-G_1223_L). Regarding the use of Ala or Leu we observed similar results.

These results indicate that the sequence and structural parameters of the TMD relevant for viral entry are distributed across the entire segment, with a stronger implication of the N-terminal portion of the TMD. It is important to note that none of the modifications included in the study significantly modified the insertion potential of the SARS-CoV-2 S protein TMD based on the predicted ΔG_app_ for their insertion (Fig 1B).

To further analyze the role of SARS-CoV-2 S protein TMD in the viral entry process we inserted alanine residues in key positions of the hydrophobic segment (Fig 2D). Inserting a residue not only increases the length of the TMD but also changes the relative orientation of the residues positioned before and after the insertion point due to the α-helical nature of this segment^30^. We first inserted an alanine between Phe_1220_ and Ile_1221_ (Ins-A_1221_) which alters the relative orientation of the aromatic membrane proximal region and the TMD. Additionally, Ins-Ala_1221_ disrupts the G_1223_xxxG_1229_ motif found in the TMD^14^, forcing a 100° rotation for Gly_1219_ relative to Gly_1223_ (Supp Fig 2A vs 2B). As shown in Fig 2D, Ins-Ala_1221_ greatly reduced the infectivity of pseudo-typed VSVΔG-GFP particles suggesting that either the relative orientation of the aromatic stretch vs the TMD, the GxxxG interaction motif, or both are important for particle entry.

Insertion of an alanine residue in positions 1226 or 1228 (Ins-A_1226_ and Ins-A_1228_) negatively affected viral entry, indicating that the relative orientation of the N-terminal and C-terminal sections of the TMD might be important for infectivity (Fig 2D). These two insertion points were selected with the work of Fu and Chou^16^ in mind, who proposed a trimerization hydrophobic zipper involving residues 1221, 1225, 1229, and 1233. Thus, the observed reduction in viral entry could also result from disrupting the oligomerization motif they described.

Insertions at positions 1230 and 1232 (Ins-A_1230_ and Ins-A_1232_) also disturb the hydrophobic trimerization motif proposed by Fu and Chou (Supp Fig. 2), but reduced particle entry to a lesser extent than Ins-A_1226_ or Ins-A_1228_ (Fig 2D). Of these two insertions, Ins-A_1230_ reduced pseudo-typed VSVΔG-GFP entry more than Ins-A_1232_, suggesting that the relative orientation of the N-terminal and C-terminal sections of the TMD is more important for the protein function than the trimerization hydrophobic zipper. Note that while the authors^16^ included leucine residues on positions 1229 and 1233 we incorporated the methionine residues found in the SARS-CoV-2 S protein consensus sequence (Uniprot: P0DTC2). Once again to ensure that the observed results are not biased by the choice of Ala residues we inserted a Trp at position 1221 (InsW_1221_) and a Met at position 1230 (InsM_1230_). Regarding the insertion of Ala or any other residue, we observed similar results (Fig 2D).

A closer look at the SARS-CoV-2 TMD reveals a highly hydrophobic surface composed of Leu_1218_, Ile_1221_, Leu_1224_, Ile_1225_, Val_1228_, and Ile_1232_ and an opposing surface rich in small residues including Gly_1219_, Ala_1222_, Gly_1223_, and Ala_1226_ which could participate in TMD-TMD interactions^15,28–31^ (Fig 3A). To analyze the importance of these regions, we first analyzed the impact of the GxxxG and the hydrophobic zipper as oligomerization motifs, we selectively mutated key residues in the SARS-CoV-2 S protein. First, we replaced Gly_1223_ with isoleucine (G_1223_I) since the equivalent substitution is sufficient to break the dimerization of Glycophorin A (GpA) TMD^31,32^. The GFP count associated with pseudo-typed VSVΔG-GFP entry revealed that the G_1223_I mutation is sufficient to reduce viral entry (Fig 3B). Substituting Gly_1219_ and Gly_1223_ with isoleucine (G_1219_I G_1223_I) further decreases pseudo-typed VSVΔG-GFP infectivity. Alternatively, we replaced Ile_1221_, Ile_1225_, or Met_1229_ from the hydrophobic surface with tyrosine (I_1221_Y, I_1225_Y, and M_1229_Y), changes that were previously suggested to modify TMD homo-oligomerization^16^. While the I_1221_Y substitution slightly reduced viral entry, the I_1225_Y, and M_1229_Y ones did not (Fig 3B).

**Figure 3.**
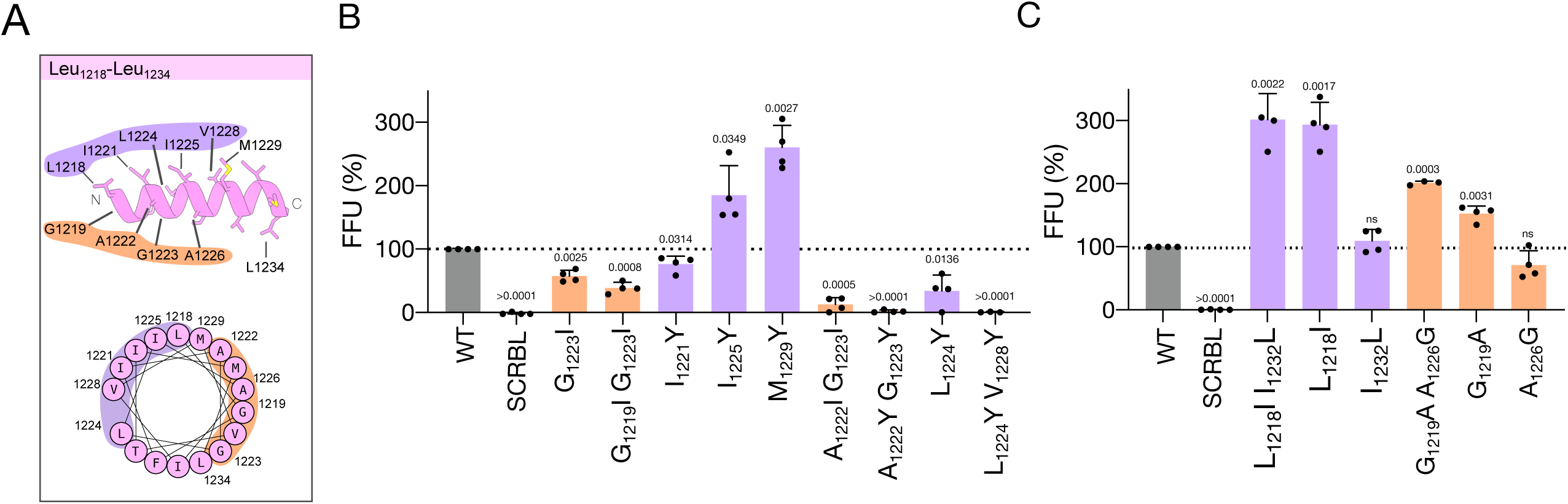
Analysis of the TMD hydrophobic and small residue-rich surfaces. **A**. Structural model of the SARS-CoV-2 S protein TMD. AlphaFold3 model of residues 1218-1234 (top) and helical wheel projection (bottom) of the same sequence. The surface of the helix containing small side chain residues is highlighted in orange, while the opposite surface containing highly hydrophobic residues is highlighted in purple. **B**. FFU from pseudotyped VSVΔG-GFP with the SARS-CoV-2 S protein bearing the following point mutations Gly_1223_ to Ile, Gly _1219_ to Ile and Gly_1223_ to Ile, Ile_1221_ to Tyr, Ile_1225_ to Tyr, Met_1229_ to Tyr, Ala_1222_ to Ile and Gly_1223_ to Ile, Ala_1222_ to Tyr, and Gly_1223_ to Tyr, Leu_1224_ to Tyr, Leu1224 to Tyr, and Val1228 to Tyr (G1223I, G1219I G1223I, I1221Y, I1225Y, M1229Y, A1222I G1223I, A1222Y G1223Y, L1224Y, L_1224_Y V_1228_Y, respectively). **C.** Analysis of the Leu_1218_ and Ile_1232_ (L_1218_I I_1232_L) and Gly_1219_ and Ala_1226_ (G_1229_A A_1226_G) residue swapping. The single mutations associated with these residue swapping Leu_1218,_ Ile_1232_, Gly_1219_, and Ala_1226_ (L_1218_I, I_1232_L, G_1229_A, and A_1226_G respectively) were also analyzed.

Then, we replaced Ala_1222_ and Gly_1223_ with isoleucine (A_1222_I G_1223_I) and measured the impact on the pseudo-typed VSVΔG-GFP infectivity. Our results revealed that the A_1222_I G_1223_I mutant is less able to promote viral entry than the previously tested G_1223_I mutant (Fig 3B). When Ala_1222_ and Gly_1223_ were replaced with tyrosine (A_1222_Y G_1223_Y), a larger and less hydrophobic amino acid, we observed a greater reduction in VSVΔG-GFP infectivity. Similarly, substituting Leu_1224_ with tyrosine (L_1224_Y) on the hydrophobic surface also reduced VSVΔG-GFP entry. Double substitution of Leu_1224_ and Val_1228_ to tyrosine (L_1224_Y V_1228_Y) diminished VSVΔG-GFP infectivity further. To further test our hypothesis we swapped Leu_1218_ and Ile_1232_ (L_1218_I I_1232_L) on one hand and Gly_1219_ and Ala_1226_ (G_1229_A A_1226_G) on the other (Fig 3C). Neither of these changes nor the single mutations associated with these residue swapping negatively affected VSVΔG-GFP entry. Therefore, according to our results, both the small residues and the hydrophobic bulky residues aligned on opposing faces of the helix (Fig 3A) are relevant for the entry process, probably because they participate in protein-protein interactions or in TMD-lipid interactions. Protein levels for all the chimeras were probed by western blotting to avoid misinterpretation of the data (Supp. Fig 1).

To incorporate the SARS-CoV-2 S protein into VSVΔG-GFP pseudo-particles it must be located at the plasma membrane. Thus, we investigated whether those events in which no VSVΔG-GFP pseudo-particle infectivity was observed were the consequence of SARS-CoV-2 S protein absence at the cell surface. We determined whether the SARS-CoV-2 S protein was located at the plasma membrane by surface-staining with an anti-RBD antibody followed by flow cytometry analysis (Fig 4). All SARS-CoV-2 S protein mutants that showed a decreased VSVΔG-GFP infectivity were included in this assay. Additionally, the WT S protein was used as a positive control while the ΔTMD was included as a negative control (Fig 4A). Our results indicate that all SARS-CoV-2 S protein mutants were present at the plasma membrane at similar levels except for Ins-A1228 (Fig 4B and Supp. Fig 3A). Thus validating the results of the VSVDG-GFP pseudo-particle assay. The absence of Ins-A1228 at the cell surface suggests that the SARS-CoV-2 S protein TMD might play a role in protein trafficking.

**Figure 4.**
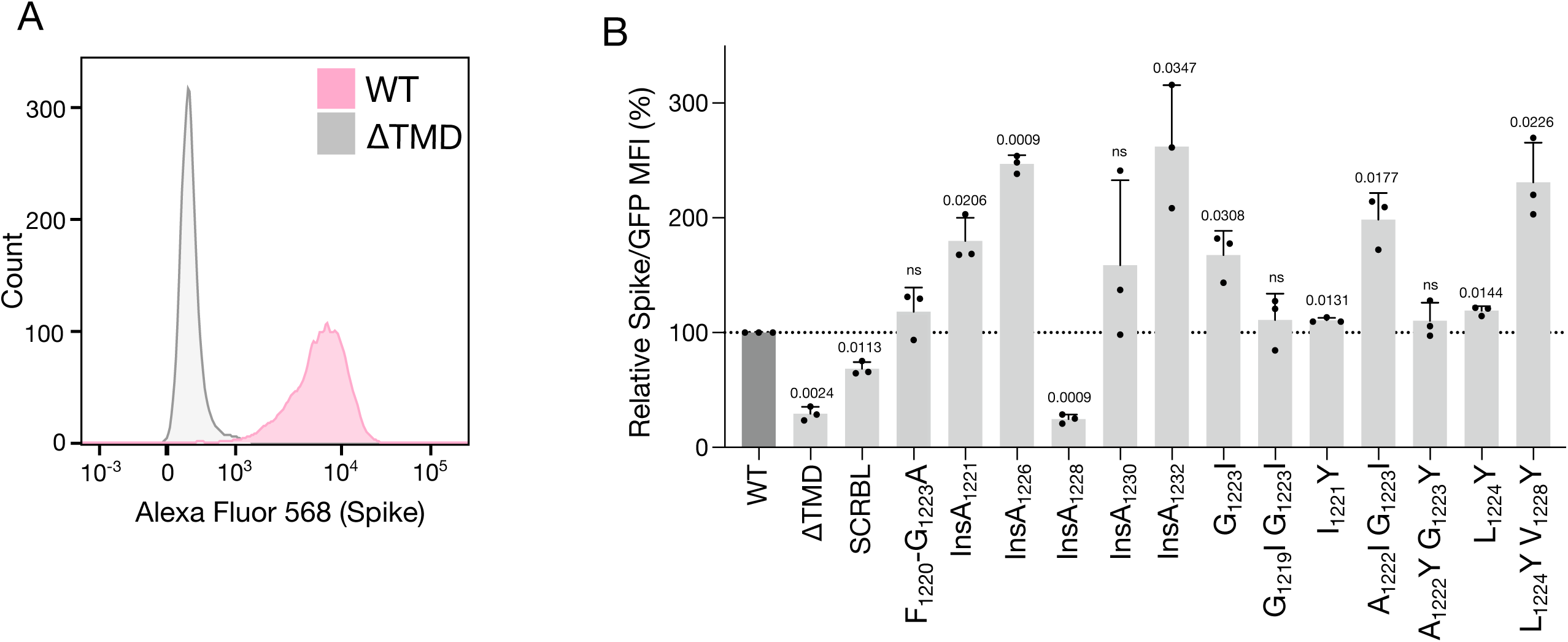
Surface expression of Spike mutants. **A.** Spike surface expression of WT and ΔTMD controls analyzed by flow cytometry. HEK293T cells were transfected with plasmids encoding the corresponding S protein alongside a constitutively GFP-expressing plasmid. After 48 hours, cells were stained with an anti-RBD antibody. Levels of the surface SARS-CoV-2 S protein (red) and GFP (green) were analyzed by flow cytometry. **B.** Relative surface expression of all Spike mutants tested in this assay. GFP was used as a transfection control. The figure shows the percentage of SARS-CoV-2 S protein vs GFP signal ratio. Samples were normalized to the SARS-CoV-2 S protein WT. The significative p-values (<0.05) for individual one-sample t-tests vs. WT are indicated above each bar.

### BiMuC assay for the analysis of membrane fusion

The role of the SARS-CoV-2 S protein TMD during the viral and cellular membrane fusion process was corroborated with a syncytia-based assay known as BiMuC^33,34^. Briefly, the Venus Fluorescent Protein (VFP) can be split into two fragments, VN and VC, neither of which is fluorescent. However, when these two fragments are fused to a pair of interacting proteins that bring them together, such as cJun and bFos, the structure of the VFP is reconstituted and the fluorescence is recovered. The VN-cJun and VC-bFos chimeras were expressed in separated cell pools together with the viral machinery required for membrane fusion. Therefore, the two chimeras would only meet in the event of membrane fusion (Fig 5A). The complete SARS-CoV-2 S WT protein was used as a positive control and reference value. The ΔTMD and SCRBL were included as negative controls (Fig 5C). Measurements of the area of the syncytia can be found in Supp. Fig 3.

**Figure 5.**
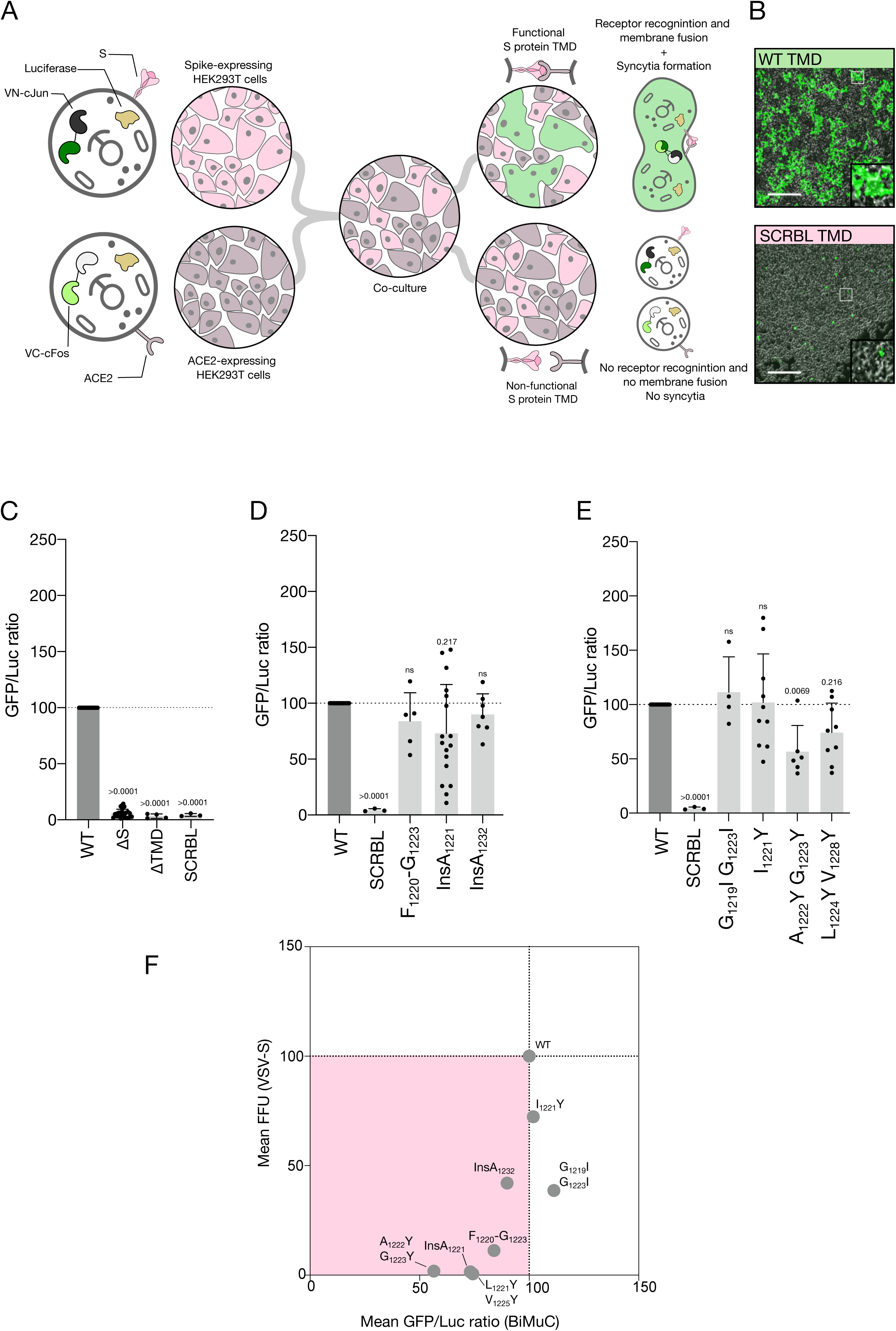
Analysis of SARS-CoV-2 S protein membrane fusion properties by BiMuC. **A**. Schematic representation of the BiMuC assay. The Venus fluorescence protein (VFP) can be split into two fragments, VN and VC respectively, neither of which is fluorescent. These two fragments were fused to cJun and bFos (VN-cJun and VC-bFos) and transfected into HEK 293T cells in separated cell pools. Cells were co-transfected with the viral machinery required for membrane fusion, the corresponding SARS-CoV-2 S protein, and the ACE2 receptor. Additionally, both cell pools were transfected with the Renilla Luciferase (luciferase) to normalize the fluorescent values. Only functional SARS-CoV-2 S proteins facilitated the S - ACE2 recognition and membrane fusion allowing VFP reconstitution and fluorescence. **B.** Representative images of the assay where cells have been transfected with the SARS-CoV-2 S protein (WT) or the SARS-CoV-2 S protein bearing a scrambled version of the TMD (SCRBL). **C-E**. The fluorescence/luciferase signal ratio (GFP/Luc ratio) was measured for the SARS-CoV-2 S protein mutants tested on this assay. Samples were normalized to the SARS-CoV-2 S protein WT. Significative p-values (<0.05) for individual one-sample t-tests are indicated above each bar. **C**. GFP/Luc ratio for the SARS-CoV-2 S protein (WT), the SARS-CoV-2 S protein where TMD was eliminated (ΔTMD), and a chimera bearing a scrambled version of the SARS-CoV-2 S protein TMD (SCRBL). As a negative control, we also included cells that did not incorporate the SARS-CoV-2 S protein (ΔS). **D**. GFP/Luc ratio for the SARS-CoV-2 S protein with substitutions of 4 amino acid stretches Phe_1220_ to Gly_1223_, and Val_1228_ to Thr_1231_, by alanines (Phe_1220_-Gly_1223_A and Val_1228_-Thr_1231_A), and the SARS-CoV-2 S protein including alanine insertions in positions 1221, 1228, and 1232 (InsA_1221_, InsA_1228_, and InsA_1232_). **E**. GFP/Luc ratio for the SARS-CoV-2 S protein bearing the following point mutations: Gly _1219_ to Ile and Gly_1223_ to Ile, Ile_1221_ to Tyr, Ile_1225_ to Tyr, Met_1229_ to Tyr, Ala_1222_ to Tyr and Gly_1223_ to Tyr, and Leu_1224_ to Tyr and Val_1228_ to Tyr (G_1219_I G_1223_I, I_1221_Y, I_1225_Y, M_1229_Y, A_1222_Y G_1223_Y, and L_1224_Y V_1228_Y, respectively). All panels include the WT values used to normalize the data (dotted line) and the SCRBL samples. **F**. Correlation of FFU measured in the VSVΔG-GFP infection assay against values of reconstituted GFP/Luc ratio in the BiMuC fusion assay. Colors show samples that have lower signals than WT values in both systems (pink background), higher in both systems (green background), or different between the two assays (white background).

Next, we selected some of the previously described mutants of the S protein and tested them using the BiMuC methodology. Substituting the Phe_1220_-Gly_1223_ helical turn with alanines (F_1220_-G_1223_A) diminished syncytia formation, though not significantly (Fig 5D). On the other hand, V_1228_-T_1231_A did not reduce syncytia formation (Fig 5D). Alanine insertions Ins-A_1221_ altered the observed syncytia-derived fluorescence (Fig 5D). However, only a small non-significant reduction in syncytia formation was observed for Ins-A_1232_. We also tested the influence of point mutations on the ability of the SARS-CoV-2 S protein to induce syncytia formation (Fig 5E). In this case, the G_1219_I G_1223_I, I_1221_Y, or the M_1229_Y substitutions did not perturb the SARS-CoV-2 S protein function. However, I_1225_Y, A_1222_Y G_1223_Y, and L_1224_Y V_1228_Y reduced the formation of syncytia. Altogether, the BiMuC-derived results correlate with the pseudo-typed VSVΔG-GFP assay^51^ (Fig 5F).

### Analysis of TMD oligomerization

Some of the modifications included in the BiMuC and the pseudo-typed VSVΔG-GFP assay could alter the previously described homo-oligomerization of the SARS-CoV-2 TMD^12,16,35^. To directly analyze the SARS-CoV-2 TMD potential oligomer, we used a bimolecular fluorescent complementation (BiFC) approach adapted to study intramembrane interactions^36–39^. Briefly, the tested TMDs were fused to a split VFP, either to its N-terminal (VN) or its C-terminal (VC). If the interaction between the tested TMDs occurs, it will bring the VN and VC ends together, facilitating the reconstitution of the VFP structure and causing fluorescence (Fig 6A). The TMD of GpA was used as a positive control and normalization value across experimental replicates. The TMD of Tomm20, a monomeric hydrophobic segment, was used as a negative control^38,40^.

**Figure 6.**
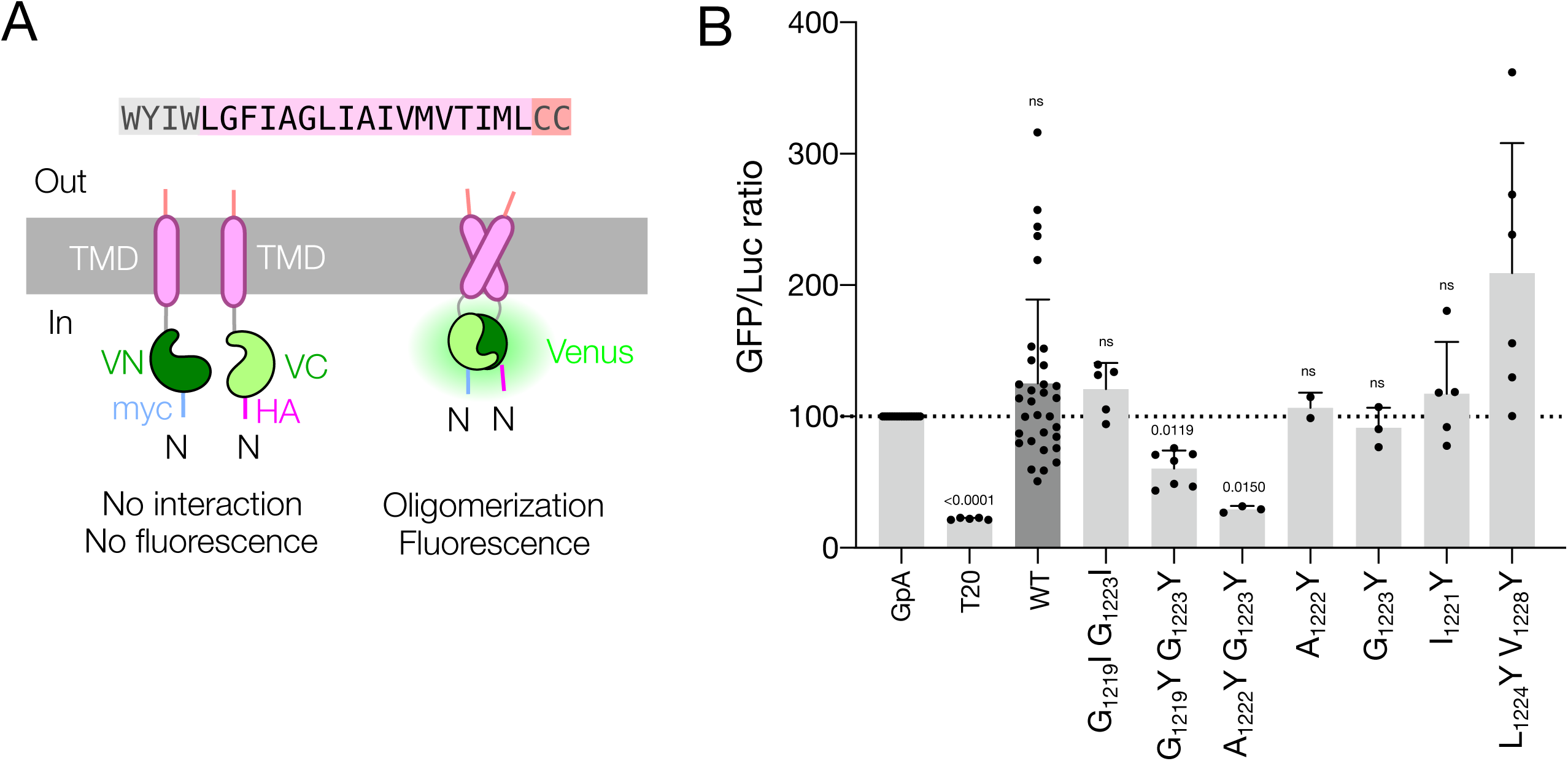
Analysis of TMD oligomerization by BiFC assay. **A**. Reconstitution of the Venus Fluorescent Protein (VFP) mediated by the SARS-CoV-2 S protein TMD oligomerization. Different S TMD mutants were fused to both the N and C terminal sections of the VFP (VN and VC, respectively) and expressed in HEK 293T cells together with the Renilla Luciferase. If the SARS-CoV-2 S protein TMD can oligomerize the VFP will be reconstituted **B.** GFP/Luc ratios measured for different chimeras bearing Glycophorin A TMD (GpA), *E.coli* Lep H2, wild-type SARS-CoV-2 S protein TMD (WT), and the SARS-CoV-2 S protein TMD with the following point mutations G_1219_I Gly_1223_ to Ile, Gly_1219_ to Tyr and Gly_1223_ to Tyr, Ala_1222_ to Tyr and Gly_1223_ to Tyr, Ala_1222_ to Tyr, Gly_1223_ to Ile, le_1221_ to Tyr, Ile_1225_ to Tyr, Met_1229_ to Tyr, and Leu_1224_ to Tyr and Val_1228_ to Tyr (G_1219_I G_1223_I, G_1219_Y G_1223_Y, A_1222_Y G_1223_Y, A_1222_Y, G_1223_I, I_1221_Y, I_1225_Y, M_1229_Y, and L_1224_Y V_1228_Y respectively). All values are normalized against GpA homo-oligomer. The significative p-values (<0.05) for individual one-sample t-tests are indicated above each bar.

Using this BiFC assay, we tested the SARS-CoV-2 TMD. Our results indicate that this TMD is sufficient for inducing oligomerization (Fig 6B). Next, we focused on the role of the surface with small residues on the oligomerization of the TMD. The G_1219_I G_1223_I substitution did not decrease the SARS-CoV-2 S protein’s TMD oligomerization potential, however, the G_1219_Y G_1223_Y did, a result that correlates with the outcome of the BiMuC assay. The A_1222_Y G_1223_Y double substitution further reduced oligomerization. Interestingly, the substitution of only Ala_1222_ or Gly_1223_ with Y (A_1222_Y and G_1223_Y), did not alter the oligomerization capabilities of the SARS-CoV-2 TMD, suggesting a synergistic effect of these two residues.

We also tested the role of the bulky hydrophobic helix interface on the SARS-CoV-2 S protein TMD on oligomerization. The single substitutions I_1221_Y, I_1225_Y, or M_1229_Y did not alter oligomerization. Furthermore, the double substitution L_1224_Y V_1228_Y, which had proven fundamental for VSVΔG-GFP entry and membrane fusion in the BiMuC assay, did not decrease the observed fluorescence, suggesting that the hydrophobic surface does not participate in the oligomerization of the SARS-CoV-2 S protein TMD.

We modeled the potential SARS-CoV-2 S protein TMD homo-trimer using TMHOP^41^. TMHOP uses Rosetta symmetric all-atom ab initio fold-and-dock simulations in an implicit membrane environment to predict low-energy conformations based on the empirical measurement of amino acid insertion propensities into *E. coli* inner membrane^42^. These conformations are clustered based on energy and structural properties (Supp Fig. 4A) and a representative model of each cluster is shown (Fig 7A). In each model, Gly_1216_, Ala_1222_, Gly_1223_, and Ala_1226_ are located on the interaction surface of the potential trimer, confirming our experimental results. We also modeled the potential TMD trimer using AlphaFold3^43^, a recent evolution of the AlphaFold2 architecture and training procedure capable of predicting the joint structure of complexes. Once again, the predicted models present, in most cases, Gly_1216_, Ala_1222_, Gly_1223_, and Ala_1226_ at the interior of a TMD homo-trimer (Fig. 7B and Supp Fig. 4). The AlphaFold3 server allowed us to model larger sequences, including the full-length S protein. When the TMD region was modeled in the context of the full-length protein accuracy was low and thus the models were discarded (Supp. Fig 5). However, the server modeled with sufficient confidence the TMD flanked by the N-terminal aromatic-rich region and the Cys-rich C-terminal end. The retrieved models for this section included, once again, the Gly_1216_, Ala_1222_, Gly_1223_, and Ala_1226_ on the internal surface of a TMD trimer (Fig. 7C and Supp Fig. 4).

**Figure 7.**
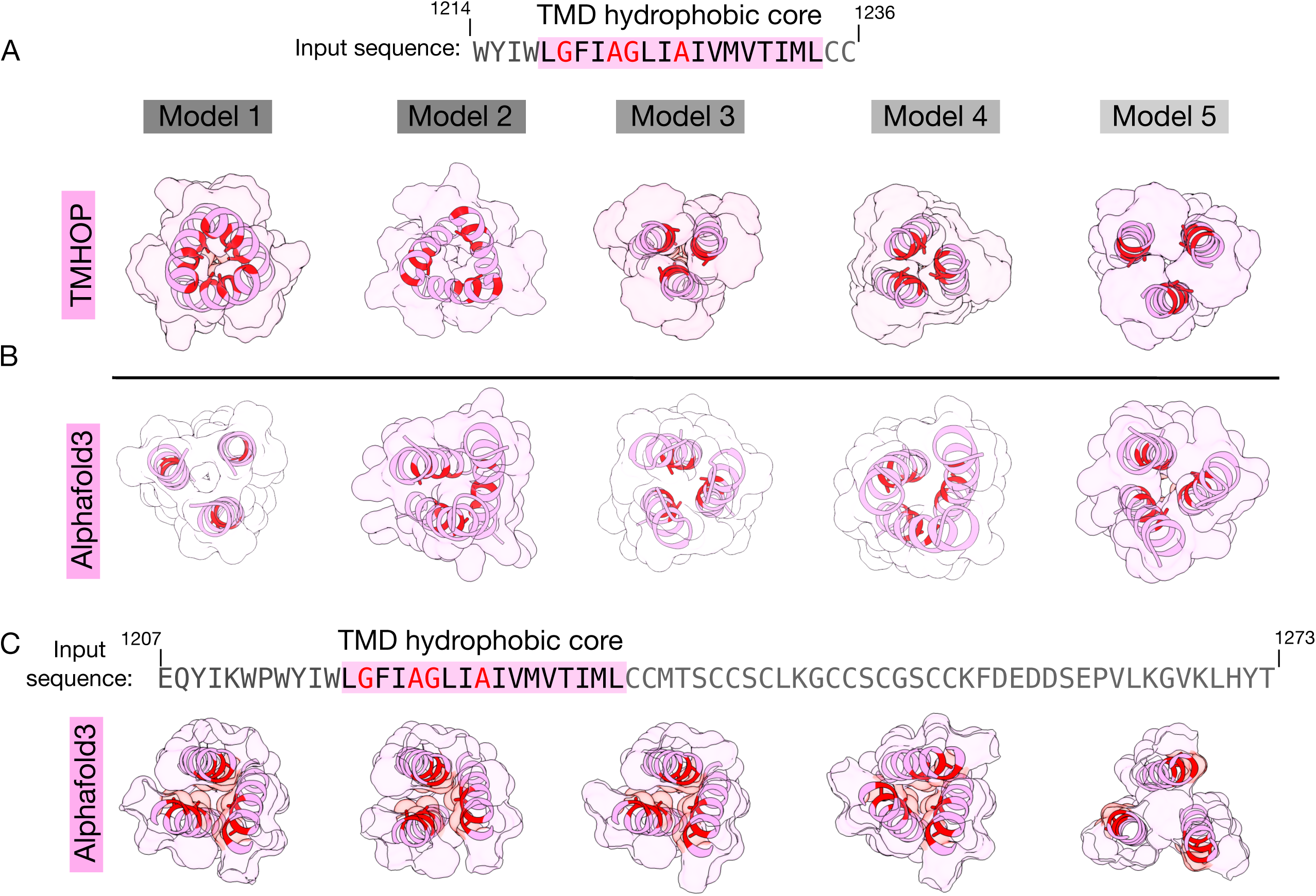
*SARS-CoV-2 S protein* TMD trimer models. **A and B.** SARS-CoV-2 S protein TMD trimer modeled by the TMHOP server (**A**) and by the AlphaFold3 server (**B**). The top five models are shown. The input sequence was _1212_WYIWLGFIAGLIAIVMVTIMLCC_1235_. Models show the TMD hydrophobic core, residues 1218-1234 shown in pink. The secondary structure as well as the surface of the peptide is shown. Gly_1216_, Ala_1222_, Gly_1223_, and Ala_1226_ are highlighted in red on the secondary structure representation. **C**. AlphaFold3 server model of the C-terminal region of the SARS-CoV-2 S protein, including the aromatic-rich section, the TMD, and the cysteine-rich domain. The five top models are shown. Models include just the TMD hydrophobic core, residues 1218-1234 shown in pink. The secondary structure as well as the surface of the TMD hydrophobic core is shown. Gly_1216_, Ala_1222_, Gly_1223_, and Ala_1226_ are highlighted in red.

## Discussion

In sum, our results indicate that the SARS-CoV-2 S protein TMD is a key region for viral entry. A variety of small changes across its sequence can render a crippled S protein. One could imagine that a passive membrane anchoring domain would accept mutations as long as the overall hydrophobicity and length are not compromised. However, a closer look at the genome of all the variants of SARS-CoV-2 reveals that no mutation whatsoever has been found in the S protein TMD, and few variations are found between SARS-CoV-2, the SARS-CoV-1, or MERS-CoV (Supp Fig 5A), while TMDs of other SARS-CoV-2 proteins present multiple mutations (Supp Fig 5B).

Within the S protein TMD, we identified at least two subdomains important for the protein’s function. One is a highly hydrophobic surface that most likely works in coordination with the lipid milieu, and the other is a surface rich in small residues (alanine and glycine) which might participate in the homo-oligomerization of the TMD. In the non-polar membrane environment, the hydrophobic effect is responsible for insertion into the membrane and does not usually contribute to the tertiary and quaternary structure^44^. Furthermore, the nature of the residues that facilitate membrane insertion impedes salt bridges and hydrogen bonds between TMDs^45^. In this context, the tertiary and quaternary structure of TMDs is dictated by a delicate balance of the remaining low energy forces, where van der Waals interactions play a crucial role^46^. The nature of van der Waals forces requires a large contact area between the associating protein segments. Amino acids with small side chains such as glycine and alanine facilitate the helix-helix contact increasing van der Waals forces. In this scenario, the inclusion of bulky residues that diminish the helix-helix contact area could break TMD-TMD interactions. Our results, together with the structural and sequence analysis, suggest that the SARS-CoV-2 S protein TMD is capable of forming homo-trimers held together by van der Waals forces. However, the structure of the full-length S protein in a post-fusion conformation did not reveal the TMD in a trimeric disposition^20^. In this structure, the TMDs surround the FPs and make contact with them through the Leu_1234_ and Phe_1120_ residues, which were not involved in either of the subdomains described in our work. Nonetheless, given that different oligomeric states coexist for the SARS-CoV-2 S protein^47^, it is conceivable that the TMDs could be found in different oligomerization states as well. Furthermore, recent simulations suggest that the SARS-CoV-2 spike TMD is inherently dynamic with different possible conformations, monomer, dimer, and trimer^35^. This conformation variety may be required for viral fusion. In the early stages of the SARS-CoV-2 S protein biogenesis, the TMD-TMD interactions responsible for the TMD trimer could facilitate the formation of a full-length protein trimer together with other motifs^8^, participate in the post-translational processing of the S protein^48^, or modulate its trafficking^49^. Next, the low-energy nature of the forces holding the TMD trimer together would allow the transition between the different oligomeric states of the protein required during the complex membrane fusion process.

Our work goes beyond understanding viral entry. Recently, TMDs and their interactions have been targeted for the modulation of protein function and the development of new therapeutic approaches^40,50^. Our results are the first step in the development of a potential new strategy for the inhibition of SARS-CoV-2 infection.

## Materials and Methods

### Plasmid constructs

Sequence encoding SARS-CoV-2 S protein (Addgene #164433) was a gift from Alejandro Balazs. All subsequent mutants were obtained with the Pfu Plus! mutagenesis kit (EurX, Gdańsk, Poland) according to the manufacturer’s instructions. For the generation of BiFC chimeric plasmids, the sequence encoding for the TMD was PCR amplified and subcloned to generate a fusion with the Nt or Ct of the Venus Fluorescent Protein (VN, VC, respectively: Addgene #27097 and #22011, a gift from Chang-Deng H). The TMD was cloned at the Ct of the VFP. These constructs were generated using In-Fusion HD cloning kit (Takara, Japan) also according to the manufacturer’s instructions. Sequences were verified by sequencing the plasmid DNA at Macrogen (Seoul, South Korea).

### Human cells and culture conditions

Human embryonic kidney (HEK293T) (from American Type Cell Collection, Manassas, VA, USA) were maintained in Dulbecco’s modified Eagle’s medium (DMEM, Gibco) supplemented with 10 % fetal bovine serum, 1 % penicillin-streptomycin (Sigma Aldrich, St. Louis, MO, USA), and 1 % MEM non-essential amino acids. Cells were maintained at 37°C and 5 % CO2 conditions. VeroE6-TMPRSS2 were obtained from the JCRB Cell Bank (catalog JCRB1819). Cells were cultured in DMEM with either 10% or 2% FBS for maintenance or infection, respectively. Selection of VeroE6-TMPRSS2 was performed using media containing 1 mg/mL G418 (Cat. no. A1720, Sigma).

### Transient DNA transfections

HEK 293T were seeded in 6, 12 or 24-well plates and grown in DMEM (Gibco) supplemented with 10 % fetal bovine serum (FBS, Gibco) the day before transfection. For the pseudo-typed VSVΔG-GFP assay, transfections were carried out with calcium phosphate. Briefly, DNA was mixed with CaCl_2_ 125 mM in HBS buffer (NaCl 140 mM, KCl 5 mM, Na_2_HPO_4_ 750 μM, dextrose 6 mM, HEPES 25 mM pH 7.05), incubated for 15 minutes and added to the HEK293 cells. For the rest of transfections, DMEM medium was mixed with plasmid DNA (up to a total of 4 μg DNA for 6-well, 2 μg for 12-well plates) and transfected into HEK293T cells adding 3 μl of PEI 1 mg/ml and 100 μl of DMEM per μg of DNA. For the surface expression analysis, 2-3 x 10^6^ HEK293T cells were transfected in 6 well-plates with the corresponding 1 μg Spike plasmid and 1 μg of a GFP encoding plasmid. In the BiMuC assay, 2 × 10^6^ HEK 293T cells in 6-well plates were transfected either with 1 μg of S and 1 μg of VN-Jun or with 1 μg ACE2 and 1 μg VC-Fos, plus 100 ng pRL-CMV.

Transfection mixtures were added dropwise onto cells and the media was changed after 24 hours. For BiFC chimeras, 1 μg of cMyc-VN-TMD and 1 μg HA-VC-TMD were transfected into 2 × 10^6^ HEK 293T cells in 12-well plates. A plasmid expressing Renilla luciferase under the CMV promoter (pRL-CMV) (50 ng) was also added to the mix to normalize the signal. For western blot (WB) analysis, 2 μg of Spike plasmid was transfected into 6-well plates containing 6 × 10^6^ HEK 293T cells.

### Pseudo-typed VSV**Δ**G-GFP assay

Pseudo-typed VSVΔG-GFP were produced by transfection of HEK 293T cells with lipofectamine 2000 (ThermoFisher Scientific) with a plasmid encoding the indicated S genes as indicated above in a 24-well plate using 750 ng DNA and 1.875 μL of lipofectamine per well. Following 24 hours, cells were infected at a multiplicity of infection of three with VSV lacking the VSV-G protein and encoding both GFP and firefly luciferase^51^ that was previously pseudotyped with the VSV-G protein to yield infectious particles. Following infection, a monoclonal antibody targeting VSV-G (a kind gift of Rafael Sanjuan, University of Valencia, Spain) was added to neutralize any remaining VSV-G bearing viruses, and infected cells were incubated for 18 hours. The supernatant was collected, clarified and frozen at -80°C. The viral titer (focus forming units [FFU] per mL) was obtained by serial dilution on VeroE6-TMPRSS2 cells in 96 well plates and counting of GFP-expressing cells using a live cell microscope following 24 hours (Incucyte SX5, Sartorius).

### Flow cytometry

Two days post-transfection, cells were washed once with PBS and detached by pipetting in 500 μl of FACS buffer (0.1% sodium azide, 0.5% BSA in PBS). Then, 200 μl of cells were collected by centrifugation at 1500 rpm for 5 min at 4°C and stained with anti-SARS-CoV-2 RBD antibody (Invitrogen, PA5-114529). After centrifugation, cells were incubated with Goat anti-Rabbit IgG (H+L) AlexaFluor 568 (ThermoFisher, A11011). Antibodies were diluted to 1:1000 in FACS buffer and 100 μl per sample were used for resuspending cells, following a 30 min incubation on ice in both cases. After three final washes with FACS buffer, surface SARS-CoV-2 spike levels were analysed using a LSRFortessa flow cytometer (BD Biosciences). Typically, 10,000 live cells were selected based on morphology and acquired. SARS-CoV-2 spike levels were assessed by gating AlexaFluor 568 (excitation at 561 nm, collection at 586/15 nm) positive cells within the gate of GFP positive cells (excitation at 488 nm, collection at 530/30 nm). Analysis was carried out using BDFACS Diva 8.0.2 and FlowJo X softwares.

### BiMuC assay

One day post-transfection, media was discarded and cells rinsed with PBS. Then, 1 mL DMEM was added to each well and one well expressing S and VN-Jun was pooled with other well expressing hACE2 and VC-Fos. Cells were counted and 500 μl containing 3 x 10^5^ cells were seeded into 4 wells of a 24-well plate. The next day, media was discarded in three of the wells, cells rinsed with PBS, collected in 100 μl PBS, and seeded into 96-well black plates for fluorescence read. Fluorescence was measured in a VictorX multi-plate reader (Perking Elmer, Waltham, MA, USA). The luminescence was measured using the Renilla Luciferase Assay kit from Sigma following the manufacturer protocol on 96-well white cell-culture plates. At least three independent experiments were done. The ratio between VFP and luciferase was registered as the relative fluorescence units (RFU) for each well. Each of those three wells was accounted as a technical replicates, and the mean of the three values was taken as a biological replicate. RFU values were then normalized against the wild-type condition. The extra well was used for imaging syncytia in a fluorescence microscope.

### Fluorescence imaging

For each condition, one well of a 24-well plate was used for imaging 24 h after co-culture of S and hACE2 expressing cells. Images were acquired with a Zeiss Axiovert 5 fluorescence microscope both with transmitted light and GFP channels with a 10X objective. Both images were overlayed and the GFP area was quantified using Fiji^52^.

### Bimolecular fluorescent complementation (BiFC) assay

Two days after transfection, PBS washed and collected for fluorescence and luciferase measurements (Victor X3 plate reader). For the Renilla luciferase readings used as signal normalization, we used the Renilla Luciferase Glow Assay Kit (Pierce, Thermofisher) according to the manufacturer’s protocol. In each experiment, the fluorescence/luminescence ratio obtained with the GpA-GpA homodimer was used for normalization. All experiments were done at least in triplicates.

### Protein expression and western blotting

Two days after transfection, media was aspirated, cells rinsed with PBS and, lysed in 300 μl of radioimmunoprecipitation assay (RIPA) buffer [150 mM NaCl, 50 mM tris-HCl (pH 8), 1% NP-40, 0.5 % sodium deoxycholate, 0.4% SDS, and 1 mM EDTA] supplemented with cOmplete EDTAfree protease inhibitors (Roche). Lysates were sonicated with three pulses of 1 s in a VCX-500 Vibra Cell sonicator (Sonics) following addition of 5X sample buffer (final concentration 62.5 mM Tris-HCl [pH 6.8], 2 % sodium dodecyl sulfate [SDS], 0.01 % bromophenol blue, 10 % glycerol and 5 % β-mercaptoethanol). Protein samples were boiled for 4 min at 95 °C, subjected to 10 % SDS-polyacrylamide gel electrophoresis (PAGE), and transferred to nitrocellulose membranes (Cytiva). Membranes were blocked for 30 min at room temperature in Tris-buffered saline supplemented with 0.05 % Tween 20 (TBS-T) containing 5% non-fat dry milk and later incubated with primary antibodies diluted in the same buffer at 4 °C overnight. Antibodies used in this study were SARS-CoV-2 Spike S1 (Invitrogen PA5-81795), rabbit anti c-Myc (Sigma PLA0001) and mouse anti-GAPDH (Santa Cruz sc-47724). Then, membranes were washed with TBS-T and incubated with goat anti-rabbit or sheep anti-mouse IgG horseradish peroxidase conjugate (Sigma DC02L or GE Healthcare NXA931, respectively) for 1 h at room temperature and washed again. All antibodies were used at a 1:10,000 dilution in TBS-T with 5 % non-fat dry milk. Detection of immunoreactive proteins was carried out using the enhanced chemiluminescence reaction (Amersham ECL Prime, Cytiva) and detected by the Image Quant LAS 4000 Mini (GE Healthcare).

### Sequence alignments

Alignments of coronavirus Spike sequences were performed using T-Coffee^53^. Aligned sequences were then exported and viewed in Jalview^54^, and residues were color-coded using ClustalX color map.

### Statistical analysis

Unpaired t-tests assuming Gaussian distribution and equal standard deviation (SD) for all conditions were applied. The p-value was two-tailed. One sample t-test was performed to compare different wild-type-normalized conditions, comparing means with a ‘hypothetical value’ of 100. Significance assessment between test samples and controls was performed using GraphPad Prism for P values < 0.05.

## Acknowledgments

We thank P. Selvi and S. Pinto for their excellent technical assistance and V. Febrer for reviewing the manuscript. This work was supported by the Generalitat Valenciana (CIPROM/2022/62) and grant PID2020-119111GB-I00, PID2023-152568NB-I00 and CNS2022-135100 by MCIN/AEI/10.13039/501100011033 and European Union NextGenerationEU/PRTR. J. O-M. has a predoctoral contract granted by the Universisty of Valencia’s “Atracció de Talent” programme (UV-INV_PREDOC 1911519). D. B. had a research contract funded by CIPROM/2022/62 project. Flow Cytometry experiments were carried out in the Cell Culture and Flow Cytometry facility at the SCSIE (University of Valencia).

---

**Supplementary Figure 1. *Expression levels of SARS-CoV-2 S protein mutants*.** Analysis by western blotting of SARS-CoV-2 S protein mutants. **A.** Expression levels of WT, ΔTMD, and SCBRL mutants. **B.** Expression levels of SARS-CoV-2 S protein mutants including alanine substitutions Phe_1220_-Gly_1223_, Leu_1224_-Ile_1227_, and Val_1228_-Thr_1231_. **C.** Expression levels of SARS-CoV-2 S protein mutants including alanine insertions in positions 1221, 1226, 1228, 1230, and 1232 (InsA_1221_, InsA_1226_, InsA_1228_, InsA_1230_, and InsA_1232_ respectively). **D.** Expression levels of SARS-CoV-2 S protein mutants including point mutations, G_1223_I, G_1219_I G_1223_I, I_1221_Y, I_1225_Y, M_1229_Y, A_1222_I G_1223_I, A_1222_Y G_1223_Y, L_1224_Y, L_1224_Y V_1228_Y. Endogenous GAPDH was used as a loading control for all experiments.

**Supplementary Figure 2. *Helical wheel diagram for SARS-CoV-2 S protein TMD hydrophobic core.* A-F.** The SARS-CoV-2 S protein TMD hydrophobic core sequence (top, residues 1218-1234), AlphaFold3 models (middle), and helical wheel representations (bottom). Residues are represented with a color code based on their predicted ΔG of insertion relative to their position within the membrane (https://dgpred.cbr.su.se/index.php?p=home).

**Supplementary Figure 3. *Area of the S protein-induced syncytia***. The area of the syncytia was calculated from fluorescent images. Each dot represents a single measurement. Bars show the are average and st. deviation of four images. The p-value was assessed by an unpaired t-test against WT condition. Values higher than 0.05 were considered non-significant (ns).

**Supplementary Figure 4. Statistical output of *SARS-CoV-2 S protein* TMD trimer models. A.** Scatterplot distribution for the S TMD homodimer modeling by TMHOP. Dots represent each of the models for the S TMD’s homotrimers. Rmsd is calculated from the lowest-energy model. The selected models are highlighted in five different colors. **B-C.** Predicted alignment error (top) for different AlphaFold3 models. The b-factor of each residue is colored by pIDDT values on the predicted structural models (bottom). Two inputs were applied: one covering just the TMD (B, residues 1214-1236) or including the N-terminal TMD flanking region up to the C-terminal end of the protein (C, residues 1207-1273).

**Supplementary Figure 5. *Statistical output of full-length SARS-CoV-2 S protein trimer models.*** Predicted alignment error (top) for different Alphafold3 models. The b-factor of each residue is colored by pIDDT values on the predicted structural models.

**Supplementary Figure 6. Mutational landscape of SARS-CoV-2 transmembrane proteome**. **A**. Sequence alignment of the SARS-CoV-2, the SARS-CoV-1, and MERS-CoV S proteins TMD. **B**. Top. Schematic representation of SARS-CoV-2 genome. ORFs that include membrane proteins are colored. The regions that encode TMDs are highlighted in gray. Bottom. Schematic representation of SARS-CoV-2 non-structural, structural, and accessory membrane proteins. Variations over the TMDs consensus sequence that are or have been circulating are indicated. Mutations in single isolates were not considered.

