## Supplementary Figure 1 for "The sequence and structural integrity of the SARS-CoV-2 Spike protein transmembrane domain is crucial for viral entry"

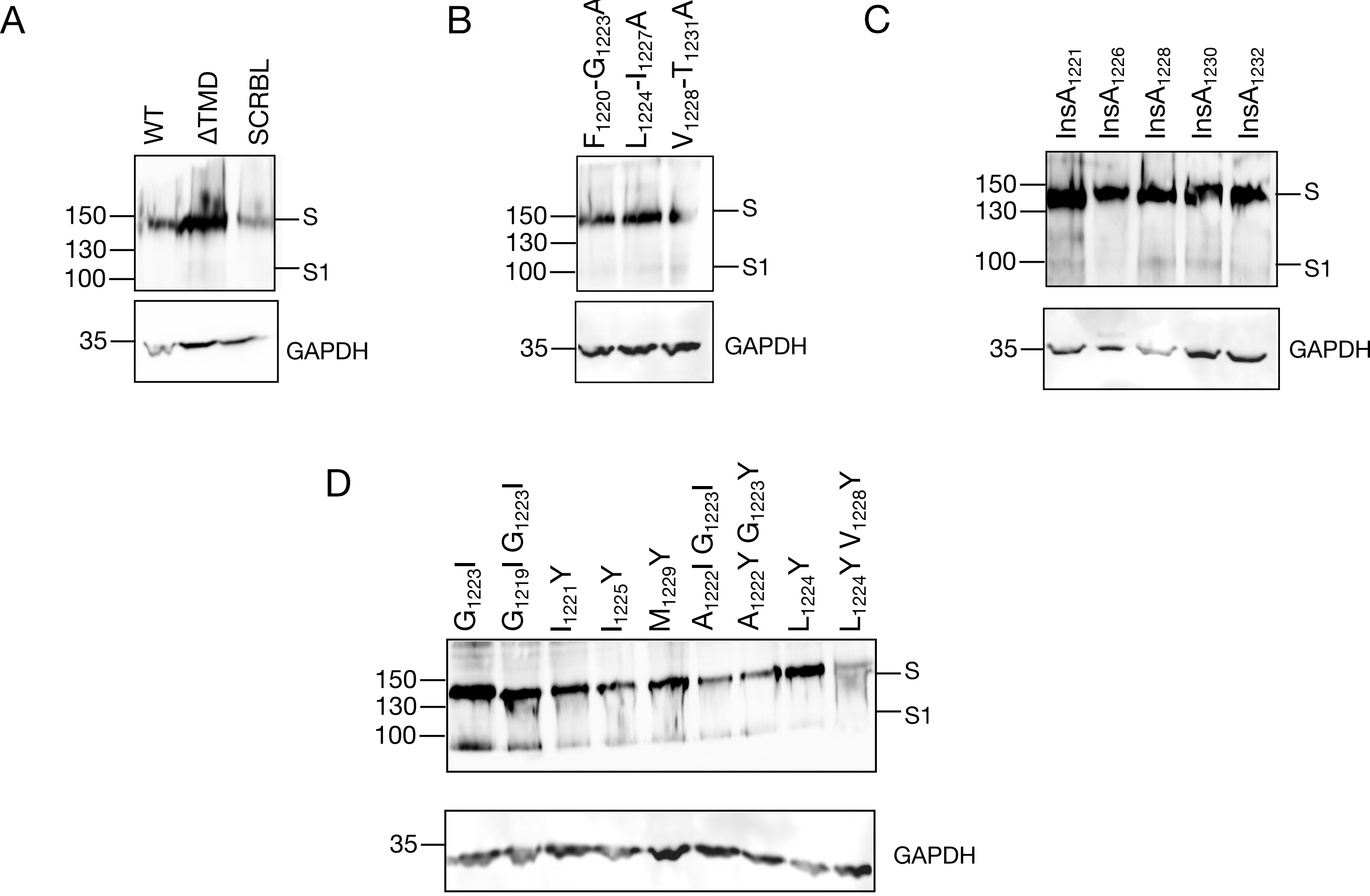


***Supplementary Figure 1. Expression levels of SARS-CoV-2 S protein variants.*** **A.** Western blot showing expression levels of WT and ΔTMD, and SCBRL variants. **B.** Expression levels of SARS-CoV-2 S protein variants including alanine substitutions Phe1220-Gly1223, Leu1224-Ile1227, and Val1228-Thr1231. **C.** Expression levels of SARS-CoV-2 S protein variants including alanine insertions in positions 1221, 1226, 1228, 1230, and 1232 (InsA1221, InsA1226, InsA1228, InsA1230, and InsA1232 respectively). **D.** Expression levels of SARS-CoV-2 S protein variants including point mutations, G1223I, G1219I G1223I, I1221Y, I1225Y, M1229Y, A1222I G1223I, A1222Y G1223Y, L1224Y, L1224Y V1228Y. Endogenous GAPDH was used as a loading control for all experiments.
