## Supplementary Figure 2 for "The sequence and structural integrity of the SARS-CoV-2 Spike protein transmembrane domain is crucial for viral entry"

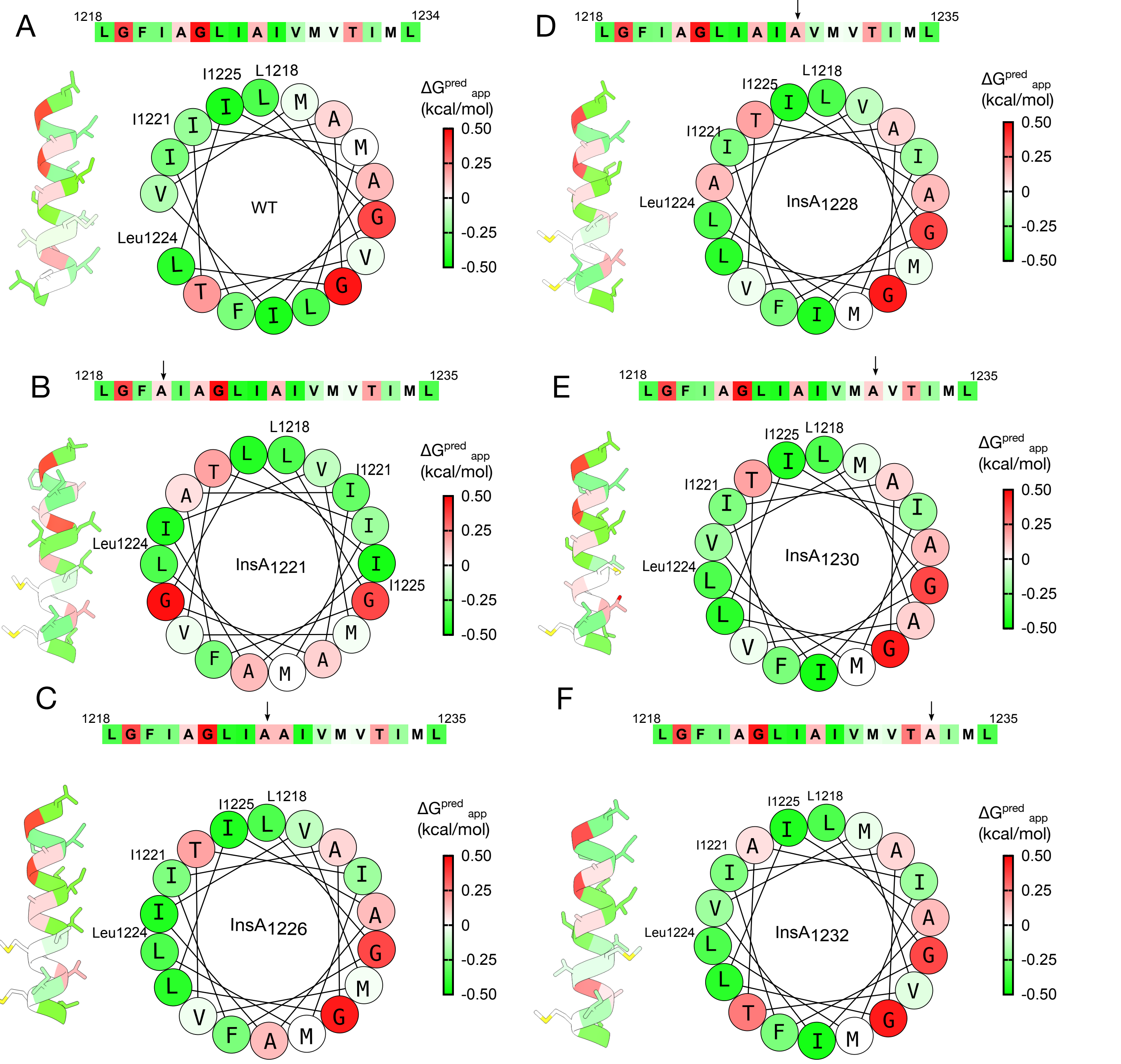


***Supplementary Figure 2. Helical wheel diagram for SARS-CoV-2 S protein TMD hydrophobic core.*** **A-F.** The SARS-CoV-2 S protein TMD hydrophobic core sequence (top, residues 1218-1234), Alphafold3 models (middle) and helical wheel representations (bottom). Residues are represented with a color code based on their predicted ΔG of insertion relative to its position within the membrane (https://dgpred.cbr.su.se/index.php?p=home).
