## Supplementary Figure 3 for "The sequence and structural integrity of the SARS-CoV-2 Spike protein transmembrane domain is crucial for viral entry"

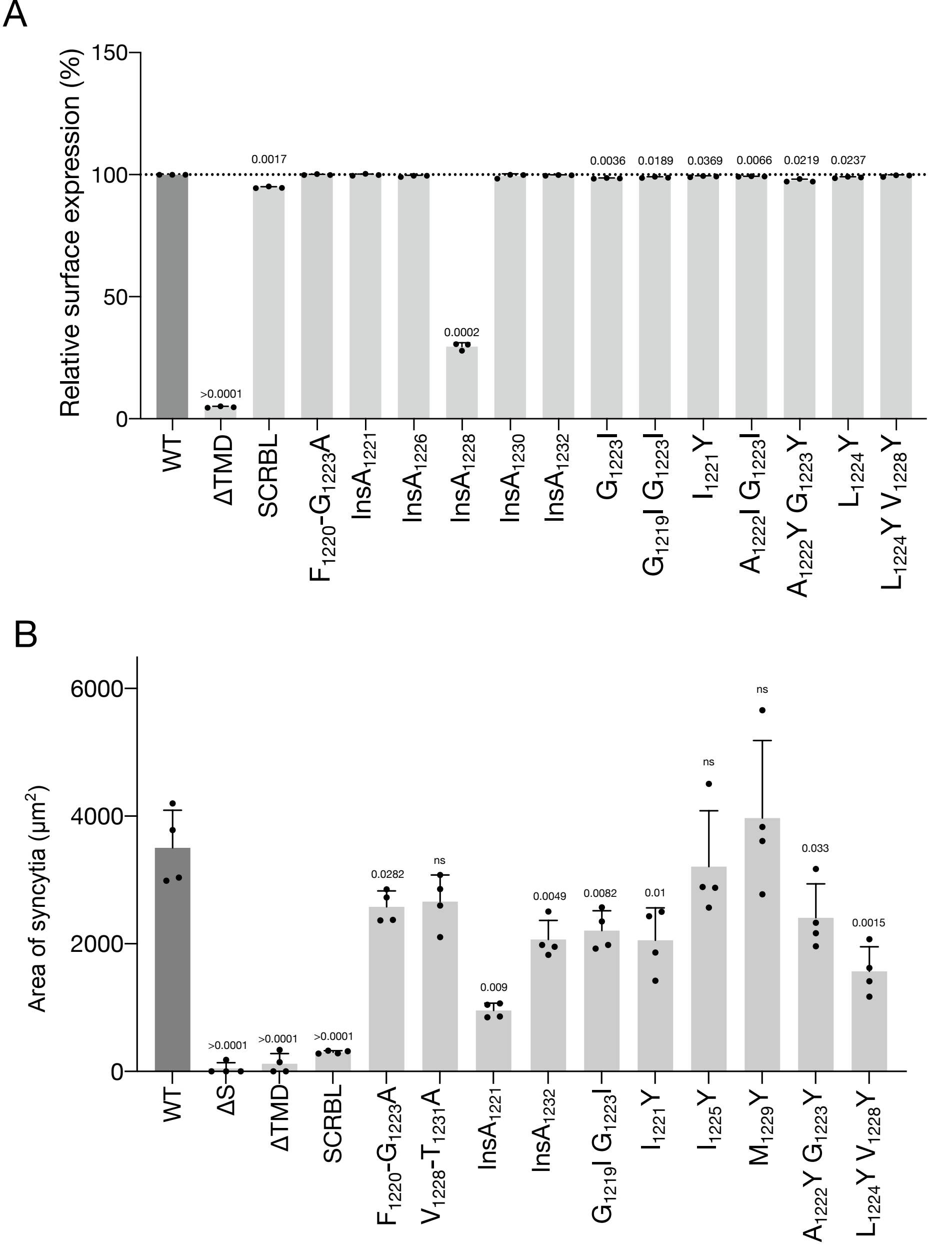


***Supplementary Figure 3. Characterization of S surface expression.*** **A.** Relative surface expression of all Spike variants tested in the flow cytometry assay. GFP was used as a transfection control. SARS-CoV-2 S protein positive cells were gated within GFP positive cells. The figure shows the percentage of double-positive events. Samples were normalized to the SARS-CoV-2 S protein WT. The significative p-values (<0.05) for individual one-sample t-tests vs. WT are indicated above each bar. **B.** Area of S induced syncytia. Total size of GFP area was calculated from images taken in a fluorescence microscope. Each dot represents a different image taken from two independent experiments. Siginificative p-values (<0.05) were assessed by an unpaired t-test against WT condition and shown above each bar.
