## Supplementary Figure 4 for "The sequence and structural integrity of the SARS-CoV-2 Spike protein transmembrane domain is crucial for viral entry"

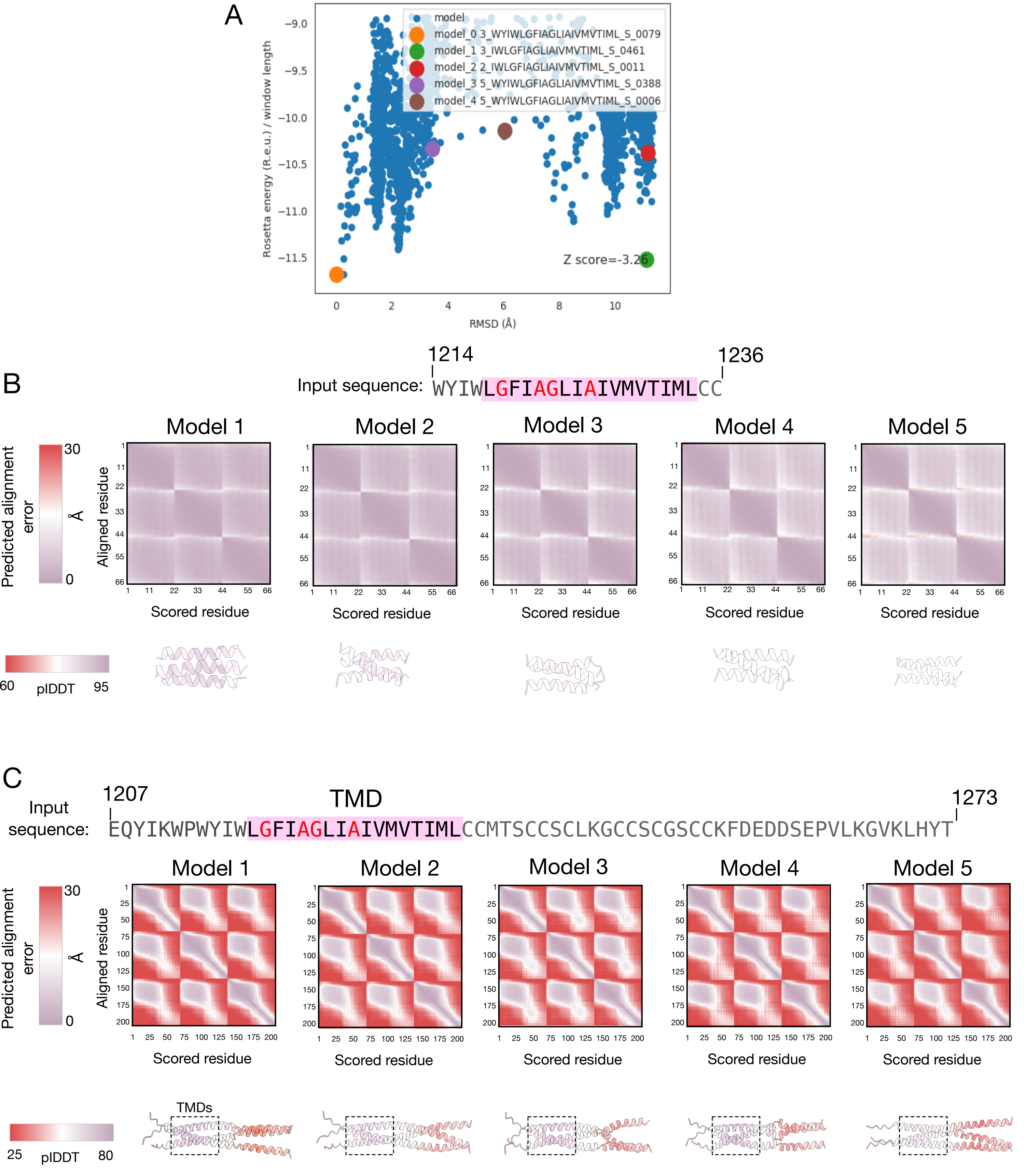


***Supplementary Figure 4. Statistical output of SARS-CoV-2 S protein TMD trimer models.*** **A.** Scatterplot distribution for the S TMD homodimer modeling by TMHOPP. Dots represent each of the models for the S TMD's homotrimers. rmsd is calculated from the lowest-energy model. The selected models are highlighted in five different colors. **B-C.** Predicted alignment error (top) for different Alphafold3 models. The b-factor of each residue is colored by pIDDT values on the predicted structural models (bottom). Two inputs were applied: one covering just the TMD (B, residues 1214-1236) or including the N-terminal TMD flanking region up to the C-terminal end of the protein (C, residues 1207-1273).
