## Supplementary Figure 5 for "The sequence and structural integrity of the SARS-CoV-2 Spike protein transmembrane domain is crucial for viral entry"

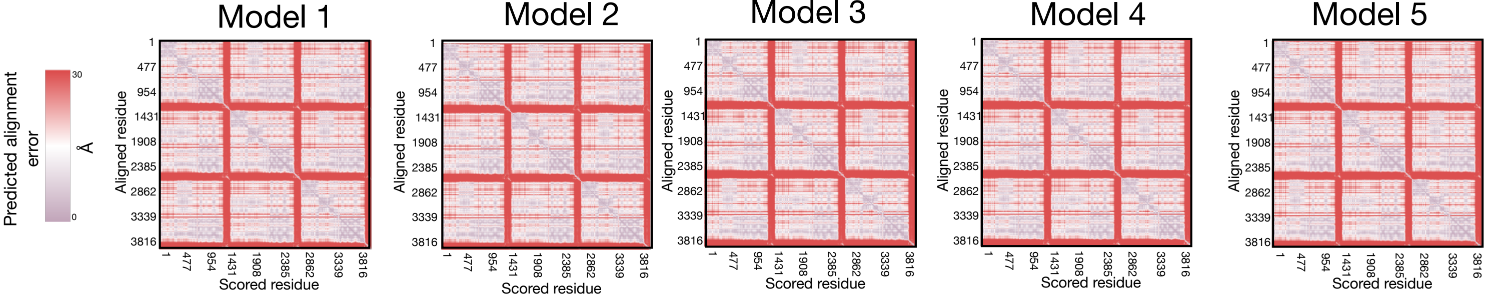


***Supplementary Figure 5. Statistical output of full-length SARS-CoV-2 S protein trimer models..*** Predicted alignment error (top) for different Alphafold3 models. The b-factor of each residue is colored by pIDDT values on the predicted structural models.
