## Supplementary Figure 6 for "The sequence and structural integrity of the SARS-CoV-2 Spike protein transmembrane domain is crucial for viral entry"

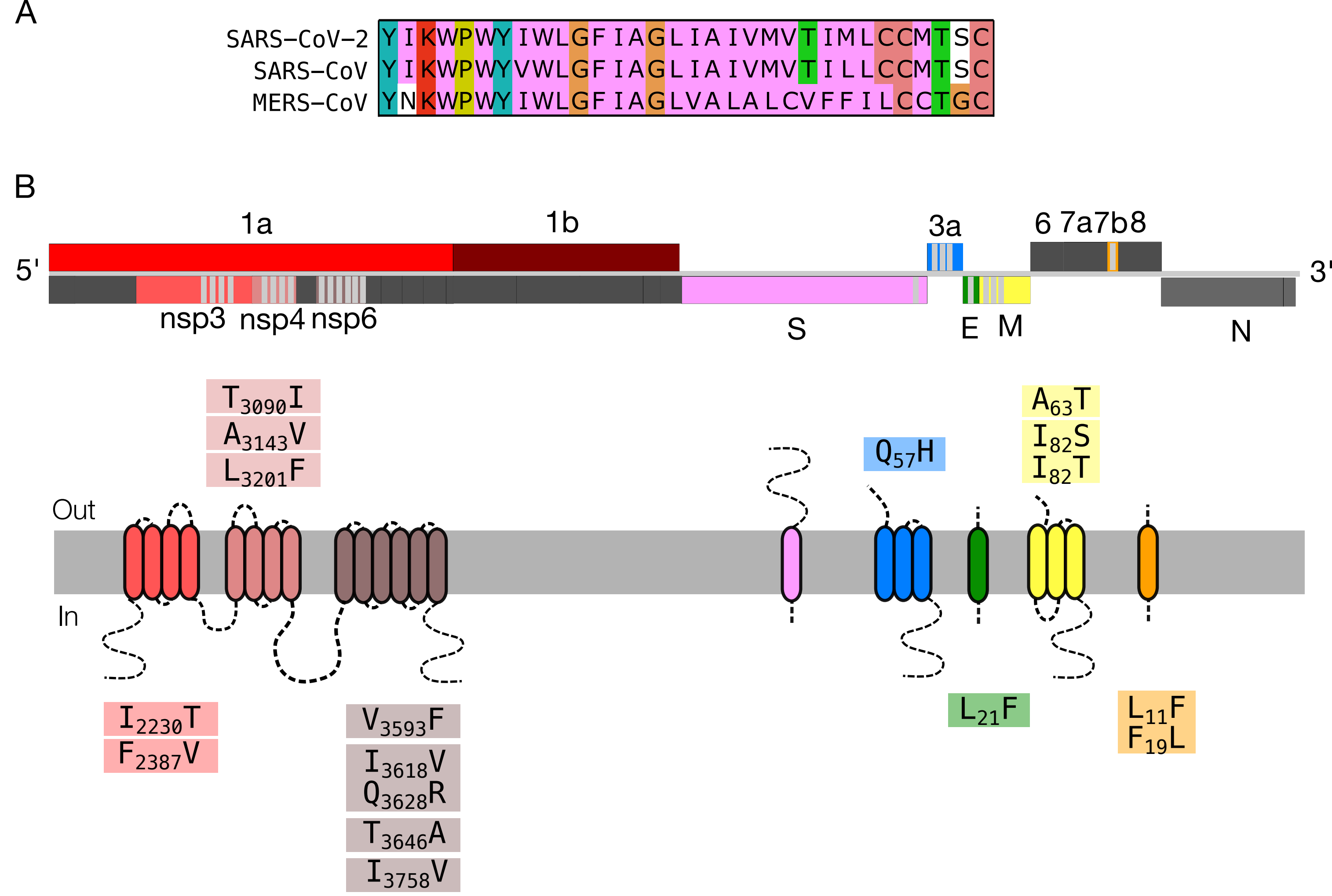


***Supplementary Figure 6. Mutational landscape of SARS-CoV-2 transmembrane proteome.*** **A.** Sequence alignment of Spike protein TMDs from SARS-CoV-2, SARS-CoV-1 and MERS-CoV. Hydrophobic residues are colored in pink, while the rest are colored by the ClustalX color scheme. **B.** (Top) Schematic representation of SARS-CoV-2 genome. ORFs that codify for membrane proteins are colored, and regions that codify for TMDs are highlighted in gray. (Bottom) Schematic representation of SARS-CoV-2 non-structural, structural and accessory membrane proteins. Accumulated mutations in different TMDs are highlighted in colors.
